# Bayesian heart-rate entropy identifies autonomic dynamics during seizure evolution: a reproducible pilot study

**DOI:** 10.64898/2026.07.30.741763

**Authors:** KR Olaciregui-Dague, FE Rosas, A Gutierrez-Gomez, R Surges, F Mormann, ML Kringelbach

## Abstract

Epileptic seizures are accompanied by profound alterations in autonomic regulation, yet the physiological information encoded in cardiac dynamics remains incompletely understood. Bayesian heart-rate (HR) entropy has recently been proposed as a probabilistic measure of cardiac dynamics, but whether it captures clinically meaningful aspects of seizure physiology—such as behavioral awareness or seizure evolution—has not been established.

We estimated Bayesian HR entropy from electrocardiographic recordings acquired during video-electroencephalographic monitoring using the BayesianAtHeart framework. Entropy-derived measures were integrated with quality-controlled clinical metadata to generate a frozen seizure-level analysis dataset, from which all subsequent analyses were performed. Associations between seizure-average Bayesian HR entropy and clinical variables were evaluated using linear mixed-effects models accounting for repeated seizures within patients, and time-resolved entropy trajectories were analyzed descriptively.

Following ECG quality control, Bayesian HR entropy was successfully estimated for 51 of 67 seizures from 10 patients; the remaining 16 seizures were excluded because ECG quality was insufficient. Forty-eight seizures with complete awareness classification comprised the primary analysis cohort. Bayesian HR entropy was not associated with ictal awareness across seizure-average analyses, mixed-effects models, or time-resolved entropy trajectories. Instead, seizure duration emerged as the strongest clinical correlate of Bayesian HR entropy, with longer seizures exhibiting progressively lower Bayesian HR entropy. Time-resolved analyses indicated that this association reflected a gradual decline in entropy during seizure evolution rather than lower Bayesian HR entropy at seizure onset. Post hoc sensitivity analyses showed that this association was not attributable to selection bias or to the number of beat-to-beat intervals available to the entropy estimator, and that duration, rather than stable between-patient differences, was the dominant source of entropy variance. Entropy was not generally reduced during seizures relative to a pre-ictal baseline; instead, the variability of the entropy trajectory declined progressively with seizure duration, more strongly than its mean.

These findings suggest that Bayesian HR entropy primarily reflects the evolving organization of autonomic regulation during seizures rather than behavioral awareness. Beyond identifying seizure duration as the strongest correlate of Bayesian HR entropy in this cohort, this study establishes a fully reproducible computational framework for Bayesian HR entropy analysis that provides a foundation for future prospective investigations of autonomic dynamics in epilepsy.

## Introduction

Epileptic seizures are accompanied by profound disturbances of autonomic function arising from dynamic interactions between cortical, subcortical and brainstem networks that regulate cardiovascular physiology.^1–5^ These autonomic responses encompass changes in heart rate, blood pressure and respiration, contribute to clinically important manifestations of seizures, and have been implicated in severe complications including sudden unexpected death in epilepsy (SUDEP).^1,6–8^ Characterizing autonomic physiology during seizures is therefore central both to understanding seizure pathophysiology and to developing objective physiological biomarkers of clinically relevant seizure characteristics.

Electrocardiography provides a practical and widely available window into autonomic regulation during seizures.^9–12^ Most previous studies have focused on heart rate or conventional heart-rate variability (HRV) metrics, including time-domain, frequency-domain and nonlinear measures.^13–15^ Although these approaches have yielded important insights, they primarily quantify the magnitude of variability over predefined time windows and may not fully capture the evolving organization of cardiac dynamics during seizure progression. Measures of physiological complexity offer a complementary perspective by characterizing the structure of autonomic regulation rather than variability alone.^16–20^

Bayesian heart-rate (HR) entropy represents one such approach. Derived using the BayesianAtHeart framework, Bayesian HR entropy estimates the uncertainty of the evolving cardiac state from beat-to-beat RR interval dynamics using probabilistic state-space modeling.^21^ By explicitly modelling cardiac dynamics as a stochastic physiological process, Bayesian HR entropy provides a quantitative measure of the complexity of cardiac dynamics that extends beyond conventional HRV metrics. While this methodology has recently been introduced and shown to capture alterations in cardiac dynamics associated with changes in global brain state, its physiological interpretation during epileptic seizures remains largely unknown.^21,22^

Several competing hypotheses are plausible. Bayesian HR entropy may reflect behavioral awareness, a clinically important feature of focal seizures that is thought to depend on the integrity of distributed cortical networks supporting consciousness.^23–25^ Because these networks are closely interconnected with the central autonomic network,^2–5^ impairment of awareness could reasonably be expected to coincide with reduced autonomic complexity. Alternatively, Bayesian HR entropy may primarily reflect seizure evolution itself, capturing progressive changes in autonomic regulation as seizures propagate through distributed neural systems.^1–5^ Distinguishing between these possibilities is essential if Bayesian HR entropy is to be interpreted as a meaningful physiological biomarker in epilepsy.

The present study therefore sought to determine which aspects of seizure physiology are most closely associated with Bayesian HR entropy. We combined Bayesian HR entropy estimation from peri-ictal electrocardiographic recordings with quality-controlled clinical seizure metadata within a fully scripted, reproducible computational workflow. Using seizure-level mixed-effects models together with time-resolved entropy analyses, we evaluated the relationship between Bayesian HR entropy, ictal awareness and seizure evolution. By identifying the clinical and physiological correlates of Bayesian HR entropy, this study aims to clarify the biological interpretation of this emerging metric and establish a reproducible analytical framework for future investigations of autonomic dynamics during epilepsy.

## Methods

### Study design and cohort

This retrospective observational study investigated the physiological correlates of Bayesian heart-rate (HR) entropy during epileptic seizures. The primary objective was to determine whether seizure-average Bayesian HR entropy differed according to ictal awareness. Secondary objectives were to characterize the temporal evolution of Bayesian HR entropy and identify clinical determinants of entropy.

Patients undergoing inpatient video-electroencephalographic (video-EEG) monitoring who experienced electrographically confirmed epileptic seizures with simultaneous electrocardiographic (ECG) recordings were included. Clinical seizure metadata were extracted from a quality-controlled seizure database containing seizure-level demographic, electroclinical and physiological variables. Seizure duration was defined as the interval between electrographic seizure onset and electrographic seizure termination as determined during routine video-EEG review.

Following quality control, 67 seizures from 10 patients were available. Bayesian HR entropy was successfully estimated for 51 seizures, comprising the entropy analysis cohort. Seizures with insufficient ECG quality for entropy estimation were excluded from entropy analyses but remained documented within the quality-control workflow. For the primary awareness analysis, seizures with unclear awareness classification were excluded, resulting in a primary analysis cohort of 48 seizures from 10 patients.

All analyses were performed using a prespecified, fully scripted computational workflow. A frozen seizure-level analysis dataset was generated before statistical modelling, and all downstream analyses were conducted exclusively from this dataset.

### Seizure classification

Ictal awareness was classified from contemporaneous clinical documentation obtained during routine video-EEG review. Seizures were assigned to one of three categories: preserved awareness, impaired awareness, or subclinical seizures with preserved awareness.^26–28^

For the primary analysis, subclinical seizures were grouped with preserved-awareness seizures because behavioral responsiveness was maintained. Seizures with unclear awareness classification were excluded from awareness-specific analyses but retained in analyses unrelated to awareness where appropriate.

### ECG preprocessing

ECG recordings (sampling rate 2048 Hz) were extracted from simultaneous video-EEG recordings using a standardized preprocessing pipeline. R-wave detection was performed once on the full continuous seizure-centered ECG segment for each seizure, using a 5−25 Hz bandpass filter (with 50 Hz notch filtering applied where mains interference was present) and an adaptive amplitude threshold (median-absolute-deviation multiplier 3.5, 85th percentile), with a minimum inter-beat interval of 0.25 s (corresponding to a maximum physiological heart rate of 240 beats min⁻^1^), to generate beat-to-beat RR interval time series; RR intervals were subsequently partitioned into baseline, peri-ictal and postictal windows for downstream analyses. Seizures were flagged for manual review or exclusion based on the burden of abnormal beat-to-beat intervals following automated detection. Seizures with inadequate ECG quality or unreliable R-wave detection were excluded before Bayesian HR entropy estimation. The identical preprocessing pipeline was applied to all seizures.

### Bayesian heart-rate entropy estimation

Bayesian HR entropy was estimated using the BayesianAtHeart framework described by Rosas et al.^21^ Bayesian state-space modelling was used to estimate the evolving probability distribution governing beat-to-beat cardiac dynamics, from which instantaneous Bayesian HR entropy was derived. This provides a principled way to study heart-rate dynamics, circumventing limitations of approaches based on frequentist statistics.^21^

Posterior inference was performed using 2,000 Gibbs sampling iterations (*N_r_* = 2000), with the first 500 iterations discarded as burn-in (*N_d_* = 500) to allow convergence of the Markov chain before entropy estimation. These sampling parameters were held constant across all analyses. Because entropy was estimated from the full sequence of beat-to-beat intervals within each seizure-centered ECG segment, the number of RR intervals available to the estimator necessarily co-varies with seizure duration; this dependency is addressed directly with a post hoc sensitivity analysis reported in the Results and discussed in the Limitations below.

For each seizure, the analysis generated a time-resolved Bayesian HR entropy trajectory, seizure-average Bayesian HR entropy, and summary heart-rate statistics.

### Construction of the frozen analysis dataset

Entropy-derived seizure metrics were merged with quality-controlled clinical seizure metadata to generate a frozen seizure-level analysis dataset used for all downstream statistical analyses. The merged dataset contained demographic, electroclinical, cardiovascular and Bayesian HR entropy variables and served as the sole input for all subsequent analyses.

### Descriptive analyses

Descriptive statistics were calculated for seizure-level demographic, electroclinical and physiological variables. Continuous variables are reported as mean ± standard deviation or median (interquartile range), as appropriate. Categorical variables are summarized as counts and percentages.

### Primary statistical analysis

The primary analysis evaluated the association between seizure-average Bayesian HR entropy and ictal awareness using linear mixed-effects regression with patient included as a random intercept to account for repeated seizures within individuals (model specification: entropy ∼ awareness + (1 | patient), fitted by maximum likelihood, with a random intercept only and no random slope). Awareness was modelled as a binary indicator (preserved/subclinical vs impaired), consistent with the seizure classification described above. Model estimates are reported with 95% confidence intervals.

### Time-resolved trajectory analyses

Time-resolved Bayesian HR entropy trajectories were aligned to electrographic seizure onset using a common peri-ictal time axis. Analyses were performed using both raw entropy values and baseline-subtracted trajectories. For each seizure, descriptive trajectory features—including baseline entropy, minimum entropy, maximum entropy, entropy range, postictal entropy and trajectory slope—were calculated. To explore associations between seizure duration and entropy dynamics, seizures were grouped into duration tertiles and mean trajectories were visualized descriptively. Owing to the limited sample size, no formal window-by-window hypothesis testing was performed.

### Exploratory determinant analyses

Associations between seizure-average Bayesian HR entropy and seizure-level clinical variables were evaluated using linear mixed-effects regression with patient included as a random intercept. Candidate explanatory variables included seizure duration, mean heart rate, awareness classification, seizure localization, seizure laterality and seizure network classification. Variables demonstrating evidence of association in exploratory analyses were subsequently entered into multivariable mixed-effects models. These exploratory comparisons were not corrected for multiple testing and are reported as hypothesis-generating rather than confirmatory. Sensitivity analyses included logarithmic transformation of seizure duration and influence analyses using Cook’s distance.

### Computational workflow and reproducibility

The complete computational workflow is summarized in **Figure 1**. The analysis pipeline comprised sequential ECG preprocessing, Bayesian HR entropy estimation, construction of a frozen seizure-level analysis dataset and downstream statistical analyses. Each analytical stage generated fixed intermediate outputs that served as the exclusive input for subsequent analyses. No downstream analysis modified previously generated datasets.

**Figure 1.**
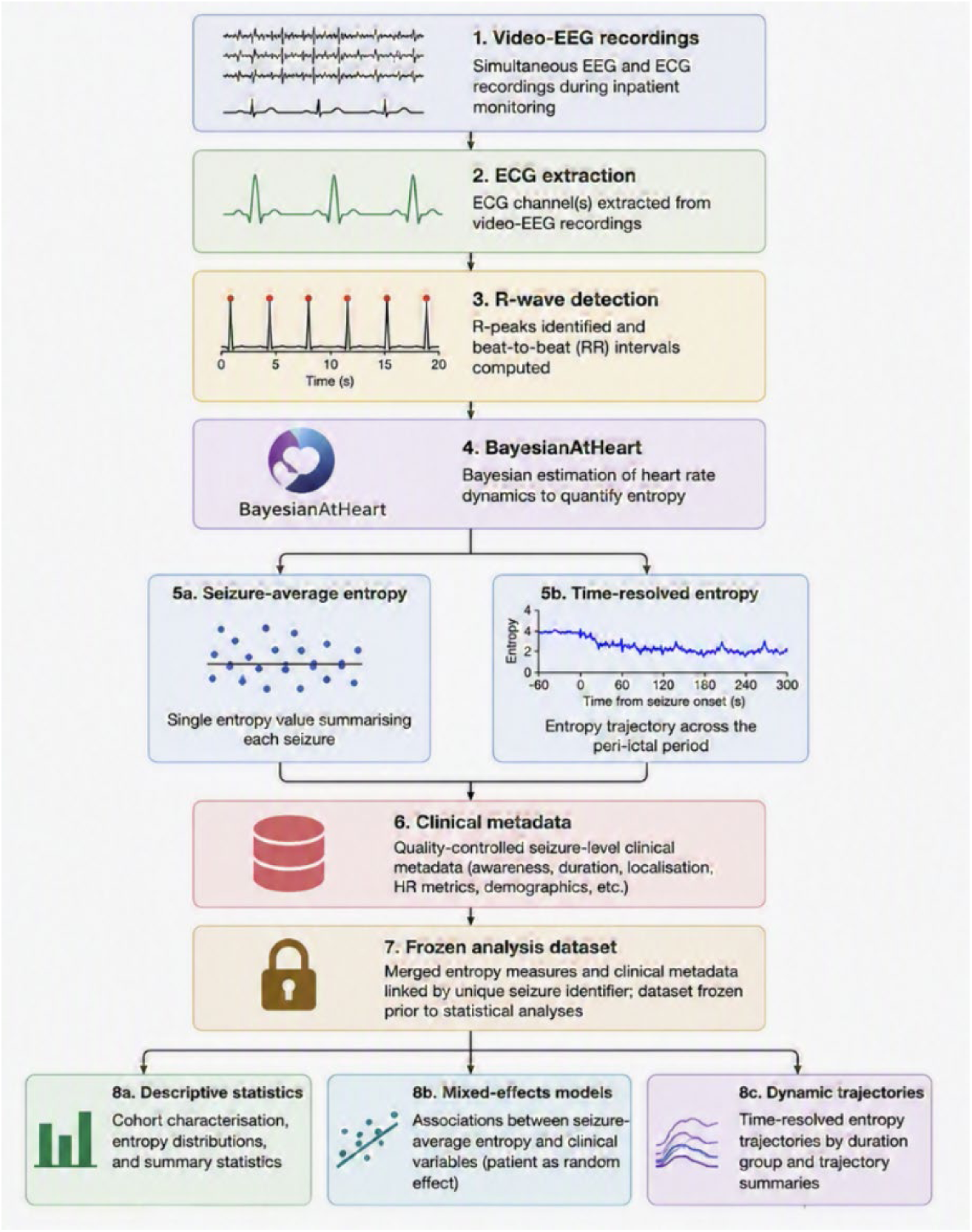
Computational workflow for Bayesian heart-rate entropy analysis. ECG recordings were extracted from simultaneous video-EEG monitoring and processed to identify R-wave peaks, yielding beat-to-beat RR interval time series. Bayesian heart-rate (HR) entropy was estimated for each seizure using the BayesianAtHeart framework, generating both seizure-average entropy estimates and time-resolved entropy trajectories. Entropy-derived measures were merged with quality-controlled clinical seizure metadata to generate a frozen seizure-level analysis dataset. All downstream descriptive analyses, mixed-effects regression models, and dynamic trajectory analyses were performed exclusively from this frozen dataset using scripted workflows. Each analysis module generated publication-ready figures, statistical tables, and accompanying source-data files without manual intervention.

The workflow comprised: (i) ECG extraction from video-EEG recordings; (ii) R-wave detection and RR interval generation; (iii) Bayesian HR entropy estimation; (iv) generation of seizure-average and time-resolved entropy measures; (v) construction of the frozen seizure-level analysis dataset through linkage with quality-controlled clinical metadata; (vi) descriptive analyses; (vii) mixed-effects regression modelling; and (viii) time-resolved trajectory analyses. Each analysis module automatically generated publication-ready tables, figures, source-data files and analysis logs.^29˒30^

### Statistical analysis

Statistical analyses were performed in MATLAB (MathWorks, Natick, MA, USA). Linear mixed-effects models were fitted using maximum likelihood estimation. Statistical significance was assessed using two-sided tests with α = 0.05. Seizure duration was z-scored (mean ≈1148 s, SD ≈162 s, based on the ECG window duration used for entropy estimation) prior to inclusion in regression models; reported duration coefficients therefore correspond to the change in Bayesian HR entropy per one standard deviation (∼162 s) increase in duration, not per second.

Given the exploratory nature of the study and the modest sample size, emphasis was placed on effect estimates, 95% confidence intervals and the consistency of findings across complementary analyses rather than dichotomous interpretation of *P* values alone.

### Post hoc sensitivity analyses

Three post hoc analyses were conducted to further probe the robustness of the duration−entropy association and the representativeness of the entropy analysis cohort. First, the awareness distribution of seizures excluded from entropy analysis because of inadequate ECG quality (n = 16) was compared with that of the entropy analysis cohort (n = 51) using Fisher’s exact test on the binary awareness classification. Second, to evaluate whether the duration−entropy association could be explained by the number of RR intervals available to the estimator scaling with seizure duration, seizure-average entropy was recomputed using only the time-resolved entropy trajectory windows falling within a fixed 0−120 s post-onset interval for every seizure, irrespective of total seizure duration, and the resulting fixed-window entropy measure was modelled against duration using the same mixed-effects specification as the primary duration model. Third, a variance-decomposition analysis quantified the proportion of total variance in seizure-average Bayesian HR entropy attributable to patient identity versus seizure duration, by comparing a random-intercept-only (null) model with a model additionally including duration as a fixed effect. These analyses were performed in Python (NumPy/SciPy) using a custom maximum-likelihood implementation of the random-intercept linear mixed model; this implementation was validated against the original MATLAB fitlme output for the primary duration model (identical coefficient, −0.012882, to six decimal places) before being applied to the sensitivity analyses. Three further post hoc analyses were conducted using the time-resolved entropy trajectories. First, to test whether Bayesian HR entropy was generally reduced during seizures relative to non-seizure periods, seizure-average entropy over post-onset windows of varying length (0−120 s, 0−300 s and 0−530 s relative to onset) was compared, within each seizure, against that seizure’s own pre-ictal baseline (window centers −530 to −300 s relative to onset), using a mixed-effects model of the paired difference with a patient-level random intercept. Second, to quantify the rate of entropy change directly, an ordinary-least-squares slope of entropy against time was estimated independently for each seizure using time-resolved entropy values from a common 0−300 s post-onset window; the resulting slopes were tested against zero and against duration and awareness using the same mixed-effects framework. As a follow-up check, this slope analysis was repeated after re-expressing each seizure’s post-onset trajectory windows as a percentage of that seizure’s own duration elapsed (0−100%) rather than absolute seconds, to test whether normalizing to relative seizure progression removed the dependence of slope on duration. Third, the standard deviation of the Bayesian HR entropy trajectory within each seizure (entropy SD, already available as a seizure-level summary statistic) was modelled against awareness and against duration using the same primary and exploratory mixed-effects specifications used for entropy mean.

### Results Study cohort

Following quality control, the study cohort comprised 67 electrographically confirmed seizures from 10 patients. Bayesian HR entropy was successfully estimated for 51 seizures (76.1%), constituting the entropy analysis cohort; the remaining 16 seizures were excluded because of insufficient ECG quality for reliable R-wave detection and entropy estimation. Three seizures with unclear awareness classification were excluded from awareness-specific analyses, yielding a primary analysis cohort of 48 seizures from 10 patients.

Across the entropy cohort, mean Bayesian HR entropy was 0.732 ± 0.030 (range 0.669−0.834), and mean heart rate during analyzed seizures was 79.5 ± 9.0 beats min⁻^1^. Within the primary cohort, 13 seizures (27.1%) were classified as impaired awareness and 35 (72.9%) as preserved awareness, including subclinical seizures with preserved awareness.

The number of seizures contributed by individual patients varied, reflecting routine clinical video-EEG monitoring. All inferential analyses accounted for repeated seizures by including patient as a random intercept in linear mixed-effects models.

### Bayesian HR entropy is not associated with ictal awareness

The primary objective of the study was to determine whether seizure-average Bayesian HR entropy differed according to ictal awareness.

Bayesian HR entropy distributions showed substantial overlap between preserved- and impaired-awareness seizures (Fig. 2). Consistent with this observation, linear mixed-effects modelling identified no association between seizure-average Bayesian HR entropy and ictal awareness (β = 0.0041, 95% CI −0.0158 to 0.0239, *P* = 0.681). This finding remained unchanged in fixed-effects analyses and following adjustment for mean heart rate.

**Figure 2.**
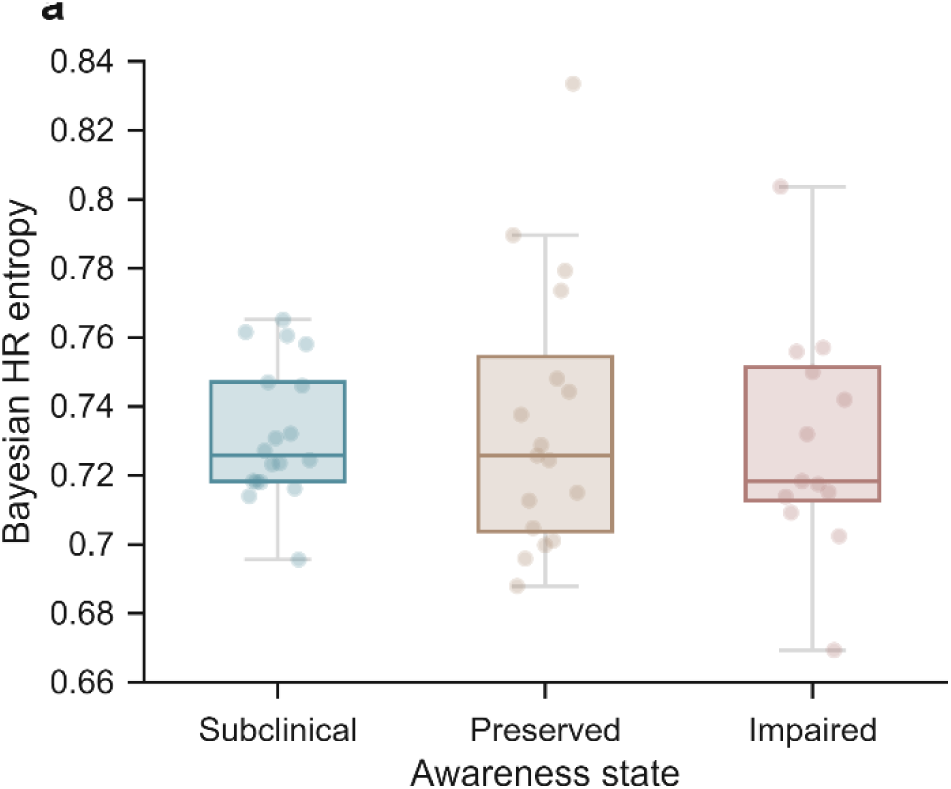
Bayesian HR entropy according to ictal awareness. Distribution of seizure-average Bayesian HR entropy across seizures with subclinical, preserved, and impaired awareness. Individual points represent individual seizures and boxplots summarize the median, interquartile range, and whiskers extending to 1.5× the interquartile range. Colors denote awareness categories. Bayesian HR entropy showed substantial overlap across awareness groups, with no consistent separation between preserved- and impaired-awareness seizures.

Time-resolved entropy trajectories likewise demonstrated no consistent separation between awareness groups (Fig. 3). Baseline-subtracted trajectories similarly failed to reveal awareness-specific entropy dynamics. Together, these findings do not support the hypothesis that Bayesian HR entropy distinguishes seizures with preserved and impaired awareness in this cohort.

**Figure 3.**
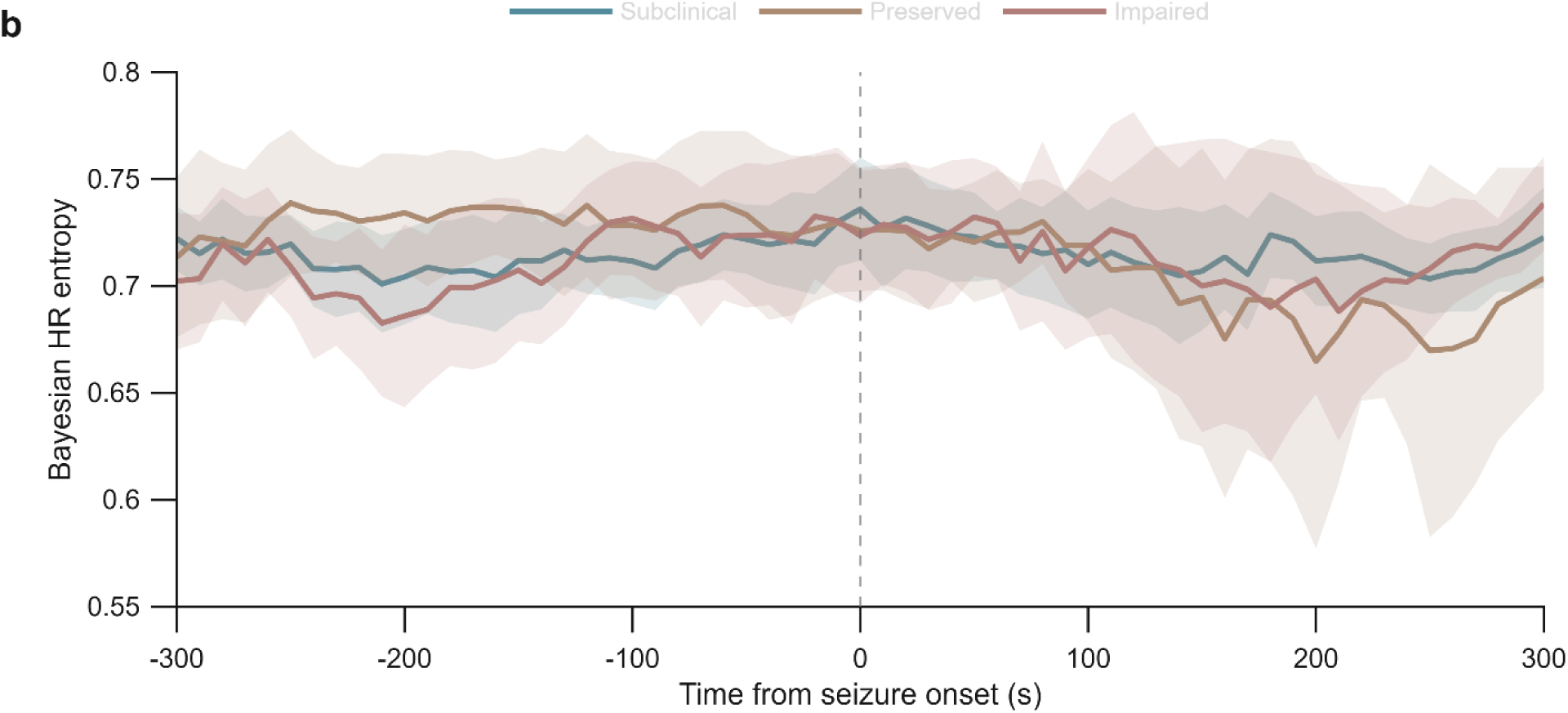
Time-resolved Bayesian HR entropy according to awareness. Mean Bayesian HR entropy aligned to electrographic seizure onset for seizures with subclinical, preserved, and impaired awareness. Solid lines represent group means and shaded regions denote 95% confidence intervals. The vertical dashed line indicates seizure onset (0 s). Entropy trajectories were broadly similar across awareness groups throughout the analyzed peri-ictal interval.

**Figure 4.**
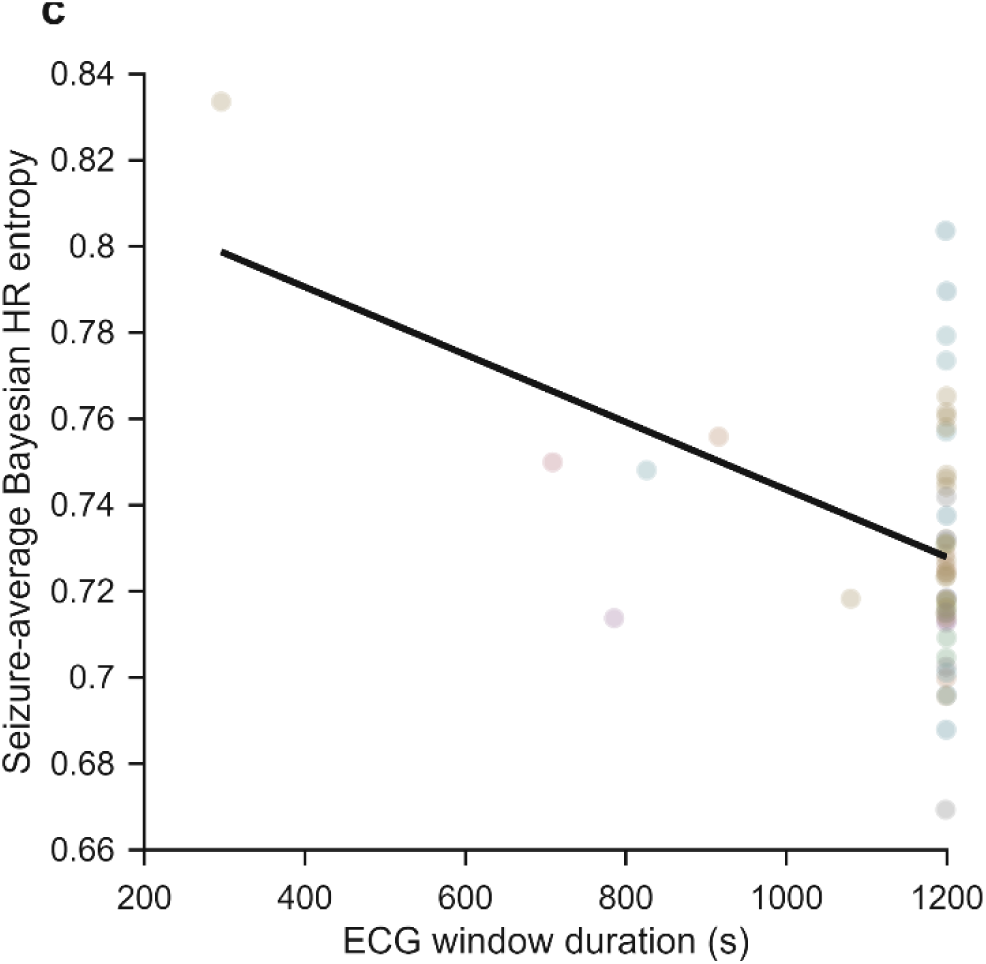
Association between seizure duration and Bayesian HR entropy. Relationship between seizure-average Bayesian HR entropy and ECG recording duration used for entropy estimation. Each point represents an individual seizure and colors denote individual patients. The solid line shows the least-squares linear fit for visualization only; statistical inference was performed using mixed-effects regression models accounting for repeated seizures within patients.

### Seizure duration is robustly associated with Bayesian HR entropy

Exploratory mixed-effects analyses were performed to identify seizure-level clinical variables associated with Bayesian HR entropy.

Among all variables examined, seizure duration showed the strongest association with seizure-average Bayesian HR entropy. Longer seizures exhibited lower Bayesian HR entropy (β = −0.0129, 95% CI −0.0204 to −0.0054, *P* = 0.0012). This association remained significant after adjustment for mean heart rate (adjusted β = −0.0124, 95% CI −0.0198 to −0.0049, *P* = 0.0017), whereas mean heart rate itself was not independently associated with entropy (*P* = 0.304). Modelling seizure duration on the logarithmic scale further strengthened the association (β = −0.0140, 95% CI −0.0214 to −0.0066, *P* = 0.00037), suggesting a nonlinear relationship.

Influence diagnostics identified a small number of seizures with relatively high leverage; however, excluding these observations did not materially alter the estimated association. Likewise, adjustment for mean heart rate produced virtually identical effect estimates, confirming that the observed relationship was not driven by differences in overall cardiac rate. The consistency of findings across unadjusted, adjusted, logarithmic and influence analyses demonstrates that the inverse association between seizure duration and Bayesian HR entropy is robust within this cohort.

By contrast, no clinically meaningful associations were observed with seizure localization, seizure laterality, seizure network classification or ictal awareness. Although beat count demonstrated a weak association with Bayesian HR entropy, it was highly correlated with seizure duration and was therefore not considered an independent explanatory variable. These analyses identify seizure duration as the strongest observed clinical correlate of Bayesian HR entropy in this cohort.

### Sensitivity analyses support the duration association and argue against major selection bias

The 16 seizures excluded from entropy analysis because of inadequate ECG quality did not differ significantly in awareness distribution from the 51 seizures comprising the entropy analysis cohort (impaired awareness: 6/16, 37.5% excluded vs 13/48, 27.1% included; Fisher’s exact *P* = 0.53), arguing against substantial awareness-related selection bias in the entropy cohort. Clinical seizure duration could not be compared between included and excluded seizures because ECG-derived duration is undefined for seizures that failed quality control.

Because Bayesian HR entropy is estimated from beat-to-beat intervals spanning the full seizure-centered ECG segment, the duration association could in principle partly reflect the number of RR intervals available to the estimator rather than a physiological effect. To evaluate this, seizure-average entropy was recomputed using only entropy trajectory windows within a fixed 0−120 s post-onset interval, held constant across all seizures irrespective of total duration. This fixed-window entropy measure remained significantly associated with (total) seizure duration (β = −0.0157, 95% CI −0.0273 to −0.0041, *P* = 0.0078), with an effect size comparable to or larger than that observed using the whole-seizure-average entropy metric (β = −0.0129). This indicates that the duration^−^entropy association is not solely an artifact of longer seizures contributing more RR intervals to the estimator, and further suggests that seizures which are ultimately longer already show lower entropy within the first two minutes after onset.

Variance-decomposition analysis indicated that patient identity explained negligible variance in seizure-average entropy in a random-intercept-only model (intraclass correlation ≈0), whereas seizure duration alone accounted for approximately 18% of total variance in entropy as a fixed effect (R^2^ = 0.181) and reduced residual (within-patient) variance by approximately 22% when added to the null model. In the duration-adjusted model, the intraclass correlation attributable to patient identity remained low (ICC ≈0.05). Together, these findings indicate that variation in Bayesian HR entropy in this cohort is explained predominantly at the seizure level by duration rather than by stable between-patient differences.

### Longer seizures exhibit progressive reductions in Bayesian HR entropy

To determine whether the association between seizure duration and Bayesian HR entropy reflected differences present at seizure onset or changes developing during seizure evolution, time-resolved entropy trajectories were stratified by duration tertiles.

Entropy trajectories were broadly similar across duration groups before seizure onset (Fig. 5). Following seizure onset, however, progressively longer seizures exhibited increasingly pronounced reductions in Bayesian HR entropy. This pattern became more apparent after subtraction of each seizure’s pre-ictal baseline. Short seizures showed a modest increase in Bayesian HR entropy relative to baseline, whereas medium- and long-duration seizures demonstrated progressively larger reductions that persisted throughout the analyzed interval.

**Figure 5.**
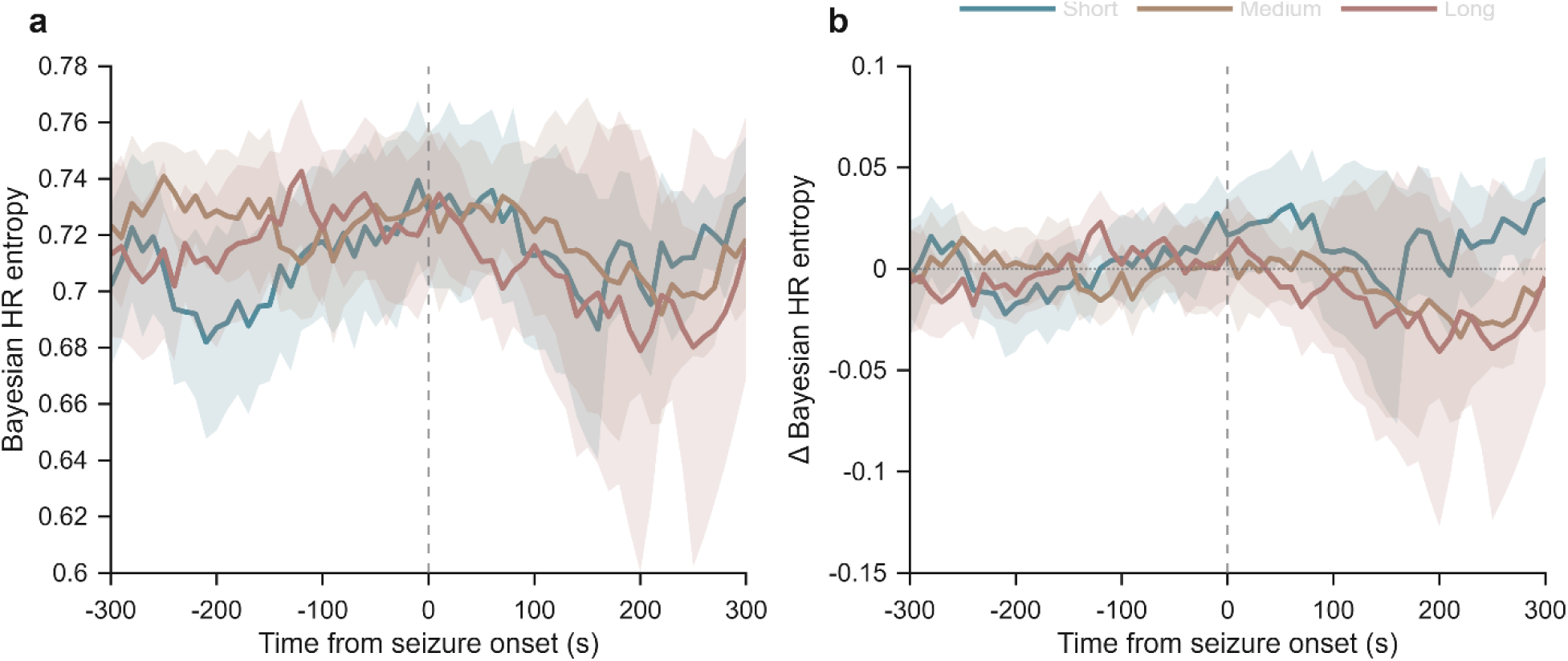
Bayesian HR entropy trajectories stratified by seizure duration. a, Mean Bayesian HR entropy trajectories for seizures grouped into duration tertiles (short, medium, and long). Solid lines indicate group means and shaded regions denote 95% confidence intervals. **b,** The same trajectories after subtraction of each seizure’s pre-ictal baseline entropy. The vertical dashed line marks electrographic seizure onset (0 s), and the horizontal dotted line indicates no change relative to baseline. Longer seizures exhibit progressively greater reductions in Bayesian HR entropy during seizure evolution.

Descriptive trajectory summaries supported these observations. The mean change from baseline to the final analysis window was positive for short seizures but negative for both medium- and long-duration seizures, with the largest reduction observed among the longest seizures.

Collectively, these findings suggest that the association between seizure duration and Bayesian HR entropy reflects progressive changes developing during seizure evolution rather than lower Bayesian HR entropy at seizure onset.

### Entropy variability, and comparison with pre-ictal baseline, further clarify the duration effect

To test whether Bayesian HR entropy is generally lower during seizures than during non-seizure periods, seizure-average entropy computed over a range of post-onset windows (0-120 s, 0-300 s and 0-530 s relative to onset) was compared against each seizure’s own pre-ictal baseline (-530 to -300 s before onset) using paired mixed-effects models. Entropy during these post-onset windows did not differ significantly from pre-ictal baseline (0-120 s: mean difference = 0.0034, 95% CI -0.016 to 0.023, *P* = 0.74; 0-300 s: mean difference = -0.0082, 95% CI -0.040 to 0.023, *P* = 0.61; 0-530 s: mean difference = -0.0029, 95% CI -0.027 to 0.021, *P* = 0.81). Bayesian HR entropy is therefore not generally depressed during seizures relative to the immediately preceding non-seizure period; instead, as shown above, entropy reductions are specific to seizures that evolve over longer periods.

To quantify the rate of entropy decline directly, seizure-specific linear regression slopes of entropy against time were estimated over a common 0-300 s post-onset window. The mean slope across the cohort was not significantly different from zero (-0.00015 entropy units per second, 95% CI - 0.00035 to 0.00006, *P* = 0.16), and slope was not associated with awareness (*P* = 0.33). Slope was, however, significantly associated with seizure duration (β = 0.000144 per SD increase in duration, *P* = 0.026): seizures that were ultimately longer showed flatter (less negative) slopes within this fixed early window, whereas seizures that were ultimately shorter showed their entire decline compressed into the same absolute interval. This indicates that entropy decline is better characterized as a function of relative seizure progression than of absolute elapsed time, and that a slope estimated over a fixed absolute window is not, by itself, a duration-independent biomarker.

To test whether normalizing time to each seizure’s own duration would remove this dependence, the slope analysis was repeated using entropy trajectory windows re-expressed as a percentage of each seizure’s own duration elapsed (0-100%) rather than absolute seconds. This normalization did not remove the duration dependence of the slope: the mean percentage-elapsed slope across the cohort was not significantly different from zero (-0.00022 entropy units per percentage point, 95% CI -0.00115 to 0.00071, *P* = 0.64), was not associated with awareness (*P* = 0.46), but remained significantly associated with duration (β = 0.0013 per SD increase in duration, *P* = 0.003) — if anything, more strongly than the absolute-time slope. This is consistent with the time-resolved trajectory shape described above: because the entropy decline in long seizures is concentrated later in the seizure course rather than distributed evenly throughout, a single whole-course linear slope, whether expressed in absolute or relative time, is diluted by an earlier flatter period and remains duration-dependent. We conclude that a simple linear slope is not a useful duration-independent summary of within-seizure entropy dynamics in this dataset; capturing this pattern would likely require a nonlinear or piecewise characterization of the trajectory, which we did not pursue further here.

Because reduced entropy could manifest as reduced trajectory variability as well as reduced mean level, we additionally examined the standard deviation of the Bayesian HR entropy trajectory within each seizure (entropy SD). Entropy SD was not associated with awareness (β = 0.0021, 95% CI -0.0049 to 0.0092, *P* = 0.55), mirroring the null result for entropy mean. However, entropy SD showed a markedly stronger association with seizure duration than entropy mean did (β = -0.0078 per SD increase in duration, *P* = 9.3 × 10^-14^, compared with *P* = 5.8 × 10^-4^ for entropy mean): longer seizures exhibited substantially less variable entropy trajectories, consistent with progressive stabilization into a more stereotyped autonomic regime as seizures evolve.

Together, these analyses indicate that the duration effect on Bayesian HR entropy reflects a progressive narrowing and stabilization of cardiac dynamics that develops over the course of a seizure, rather than a uniform reduction present from seizure onset, and suggest that reduced entropy variability may be an even more sensitive marker of this process than reduced mean entropy.

## Discussion

Bayesian heart-rate (HR) entropy has recently emerged as a principled measure of the complexity of cardiac dynamics, yet its physiological significance during epileptic seizures has remained unclear. In this pilot study, we combined Bayesian HR entropy estimation with a fully reproducible computational analysis pipeline to investigate the relationship between cardiac entropy and seizure physiology. Although Bayesian HR entropy was not associated with behavioral awareness, we identified a robust and consistent relationship with seizure duration. Time-resolved analyses further demonstrated that this association was driven not by lower entropy at seizure onset, but by a progressive decline in entropy as seizures evolved. Collectively, these findings suggest that Bayesian HR entropy primarily reflects the dynamic evolution of autonomic regulation during seizures rather than behavioral awareness itself.

The absence of an association between entropy and awareness is an important finding. Loss of awareness is among the most clinically relevant manifestations of focal epilepsy and has frequently been linked to widespread cortical and subcortical network dysfunction.^23–25^ We therefore hypothesized that disruption of consciousness might be accompanied by a measurable reduction in autonomic complexity. Instead, seizure-average entropy estimates, mixed-effects models and time-resolved trajectories consistently demonstrated substantial overlap between seizures with preserved and impaired awareness. While the present cohort was modest in size, the agreement across complementary analytical approaches suggests that any effect of awareness on Bayesian HR entropy is likely to be considerably smaller than that of seizure duration. These findings therefore argue against behavioral awareness as the principal physiological determinant of cardiac entropy during seizures.

Negative biomarker studies are valuable when they constrain competing biological hypotheses. Our findings suggest that behavioral state and autonomic entropy are at least partially dissociable, indicating that alterations in cardiac entropy cannot be inferred directly from impairment of consciousness. Rather than diminishing the value of Bayesian HR entropy, this negative result helps define the physiological domain in which the metric is likely to be informative.

In contrast, seizure duration emerged as the dominant determinant of Bayesian HR entropy across all analyses. Longer seizures consistently exhibited lower entropy, and this relationship remained stable following adjustment for mean heart rate, across multiple model specifications, and after influence analyses. Importantly, time-resolved trajectory analyses demonstrated that long seizures did not begin with markedly lower entropy than short seizures. Instead, entropy progressively declined relative to each seizure’s own pre-ictal baseline as seizure duration increased. Taken together, the static and dynamic analyses therefore indicate that Bayesian HR entropy is coupled more closely to seizure evolution than to behavioral state. Two post hoc analyses reinforced this interpretation: the duration association persisted when entropy was recomputed from a fixed post-onset window common to all seizures, arguing against a purely estimator-driven artifact of longer seizures contributing more beat-to-beat intervals; and variance-decomposition analysis indicated that duration, rather than stable between-patient differences, was the dominant source of variance in entropy, consistent with a seizure-level rather than trait-level process.

A more general, complexity-theoretic consideration may help build intuition for why seizure duration, rather than awareness, tracks Bayesian HR entropy. Work on entropy-rate-based complexity measures in neural signals has argued that such measures admit two complementary readings: as an index of the diversity of trajectories a system explores over time, and as an index of the system’s intrinsic unpredictability, or prediction error, even given complete knowledge of its own past.^39^ Applying this framework by analogy to the present findings, seizures are characterized by increasingly stereotyped, rhythmic, repetitive electrographic discharges as ictal activity propagates and entrains distributed networks. Such stereotypy would, a priori, be expected to restrict the diversity of states available to downstream physiological systems coupled to this activity and to render their moment-to-moment dynamics more predictable, and thus to lower measured entropy, independent of any specific relationship to behavioral awareness. This offers an intuitive rationale for why seizure duration, which indexes cumulative exposure to this increasingly stereotyped regime, would be expected to track progressively declining cardiac entropy, and is complementary to the network-recruitment account proposed below. Notably, direct comparison of ictal and pre-ictal entropy did not show a general reduction in entropy from the non-seizure baseline; rather, the reduction was specific to seizures that persisted longer, and was considerably more pronounced in entropy variability (SD) than in entropy mean. This is consistent with the stereotypy account above in its more precise form: it is prolonged exposure to repetitive ictal dynamics, rather than the mere presence of seizure activity, that is expected to progressively narrow the diversity of explored states and stabilize (i.e., reduce the variability of) cardiac dynamics.

One possible mechanistic interpretation is that Bayesian HR entropy reflects the progressive recruitment of distributed central autonomic networks during seizure propagation (Fig. 6). The central autonomic network—including the insular cortex, anterior cingulate cortex, amygdala, hypothalamus and brainstem autonomic nuclei—coordinates cardiovascular regulation through highly interconnected cortical and subcortical pathways.^2–5^ As seizures evolve, increasing recruitment of these structures may progressively constrain autonomic flexibility, resulting in more stereotyped cardiac dynamics and consequently lower Bayesian HR entropy. This interpretation is broadly consistent with previous work demonstrating widespread autonomic activation during epileptic seizures, seizure-related alterations in cardiac rhythm and heart-rate variability, and the close anatomical relationship between cortical networks supporting consciousness and central autonomic control.^9–12,23–25,31–37^ Under this model, declining entropy would not simply represent reduced variability in heart-rate dynamics, but a progressive loss of flexibility within distributed autonomic control systems as seizure activity evolves. Although biologically plausible, this interpretation remains hypothetical and will require direct evaluation using multimodal studies integrating electrophysiology, autonomic physiology and structural or functional neuroimaging.

**Figure 6.**
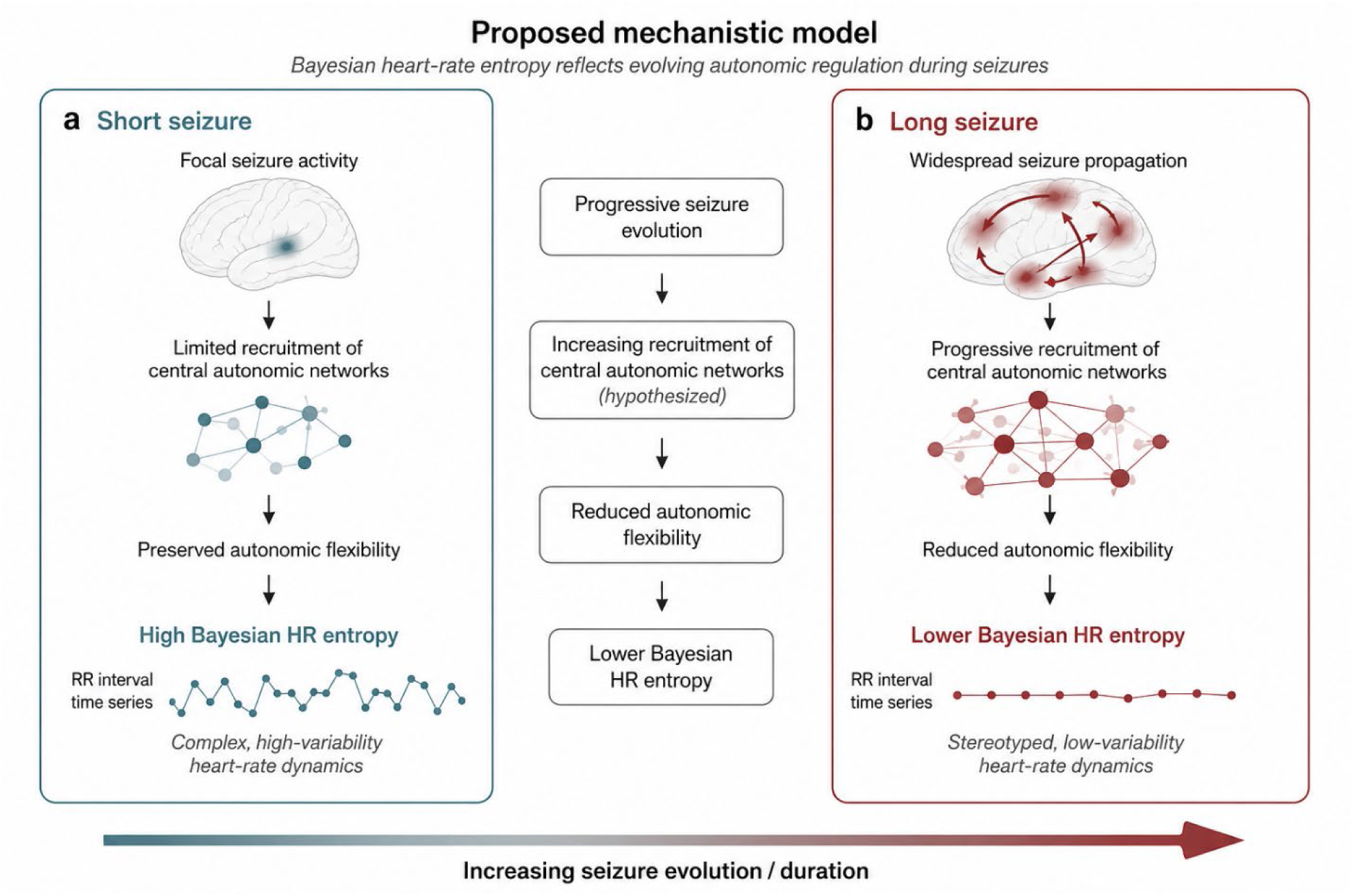
Proposed mechanistic model linking Bayesian heart-rate entropy to seizure evolution. a, During shorter seizures, relatively limited recruitment of central autonomic networks is hypothesized to preserve autonomic flexibility, resulting in higher Bayesian heart-rate (HR) entropy and more complex beat-to-beat cardiac dynamics. **b,** As seizures evolve and propagate through distributed neural networks, progressively greater recruitment of central autonomic networks is hypothesized to constrain autonomic flexibility, leading to lower Bayesian HR entropy and increasingly stereotyped cardiac dynamics. The central panel summarizes the proposed sequence linking seizure evolution to progressive reductions in Bayesian HR entropy. This conceptual model is derived from the present findings and existing knowledge of central autonomic network organization and is intended as a hypothesis to guide future mechanistic studies rather than as a direct inference from the data.

Our findings also distinguish Bayesian HR entropy from conventional autonomic biomarkers. Heart rate, classical heart-rate variability metrics and electrodermal activity have all been investigated as markers of seizure severity and behavioral impairment,^9–20,31–37^ but each captures only selected aspects of autonomic physiology. Bayesian HR entropy instead quantifies the evolving complexity of cardiac dynamics themselves.^21^ Previous work demonstrated that Bayesian HR entropy is sensitive to changes in global brain state during psychedelic drug administration.^22^ The observation that entropy tracked seizure evolution rather than awareness in the present study therefore suggests that Bayesian HR entropy provides complementary physiological information regarding the organization of autonomic regulation during seizures. Rather than functioning as a diagnostic marker of impaired awareness, Bayesian HR entropy may be better viewed as a physiological descriptor of seizure evolution.

Beyond the biological findings, this study establishes a fully reproducible computational framework for Bayesian HR entropy analysis. All preprocessing, entropy estimation, statistical analyses and figure generation were performed using scripted workflows operating on a frozen seizure-level dataset, with each analytical stage generating publication-ready tables, figures and accompanying source-data files. Although computational reproducibility does not itself validate a biomarker, it facilitates transparent evaluation, improves analytical robustness and provides a framework that can be directly applied to independent datasets and future prospective studies.^29,30^

Several limitations should be acknowledged. First, this was a retrospective pilot study comprising 51 seizures from 10 patients, limiting statistical power for subgroup analyses and reducing the precision of patient-level variance estimates. The unequal distribution of seizures across patients reflects routine clinical practice but also reinforces the importance of accounting for repeated observations using mixed-effects models. Second, awareness classification relied on clinical assessment during video-EEG monitoring and therefore remains subject to the inherent limitations of behavioral evaluation. Third, entropy estimation was restricted to seizures with ECG recordings of sufficient quality for reliable beat detection, potentially introducing selection bias; the excluded and included seizures did not differ significantly in awareness distribution (Fisher’s exact P = 0.53), though a comparable comparison of seizure duration was not possible because ECG-derived duration is undefined for excluded seizures. Fourth, because R-wave detection and Bayesian HR entropy estimation were performed over the full seizure-centered ECG segment, the number of beat-to-beat intervals contributing to each seizure’s entropy estimate necessarily scales with seizure duration. A post hoc fixed-window sensitivity analysis, in which seizure-average entropy was recomputed from a common 0^−^120 s post-onset interval for every seizure regardless of total duration, showed that the duration association persisted (and was, if anything, numerically larger) under this fixed-length estimation window, arguing against a simple estimator-sequence-length artifact as the explanation for the observed association; nonetheless, this analysis reused the existing sliding-window entropy trajectories rather than re-running the Bayesian estimator from raw beat-to-beat data on truncated recordings, and a full re-estimation from raw RR intervals would provide a more definitive test. A complementary variance-decomposition analysis indicated that seizure duration, not stable between-patient differences, is the dominant source of variance in Bayesian HR entropy in this cohort. Finally, although Bayesian HR entropy was consistently associated with seizure duration, the observational design precludes conclusions regarding causality or the underlying neural mechanisms.

These limitations also define the next stage of investigation. The present study establishes feasibility and provides effect-size estimates for Bayesian HR entropy in epilepsy, but confirmation will require substantially larger prospective cohorts with balanced patient representation and predefined trajectory analyses. Integration with quantitative measures of seizure propagation, autonomic physiology and multimodal neuroimaging may clarify the biological mechanisms responsible for the progressive entropy reductions observed during prolonged seizures. Such studies will also determine whether entropy dynamics predict clinically meaningful outcomes, including seizure termination, secondary generalization or postictal recovery.

In conclusion, Bayesian HR entropy was not associated with ictal awareness in this pilot cohort. Instead, Bayesian HR entropy decreased progressively during longer seizures, indicating that it reflects the dynamic evolution of autonomic regulation rather than behavioral awareness. These findings identify seizure duration as the principal physiological determinant of cardiac entropy in this dataset and provide a reproducible analytical framework for future investigations of autonomic dynamics during epileptic seizures.

## Data and code availability

All preprocessing, entropy estimation, statistical analysis and figure-generation code used in this study is available at Github upon request. The frozen seizure-level analysis dataset underlying the statistical results reported here is available from the corresponding author upon reasonable request, subject to institutional and data-protection approvals governing the underlying clinical video-EEG and ECG recordings.

