## Supplementary Table 1 for "Bayesian heart-rate entropy identifies autonomic dynamics during seizure evolution: a reproducible pilot study"

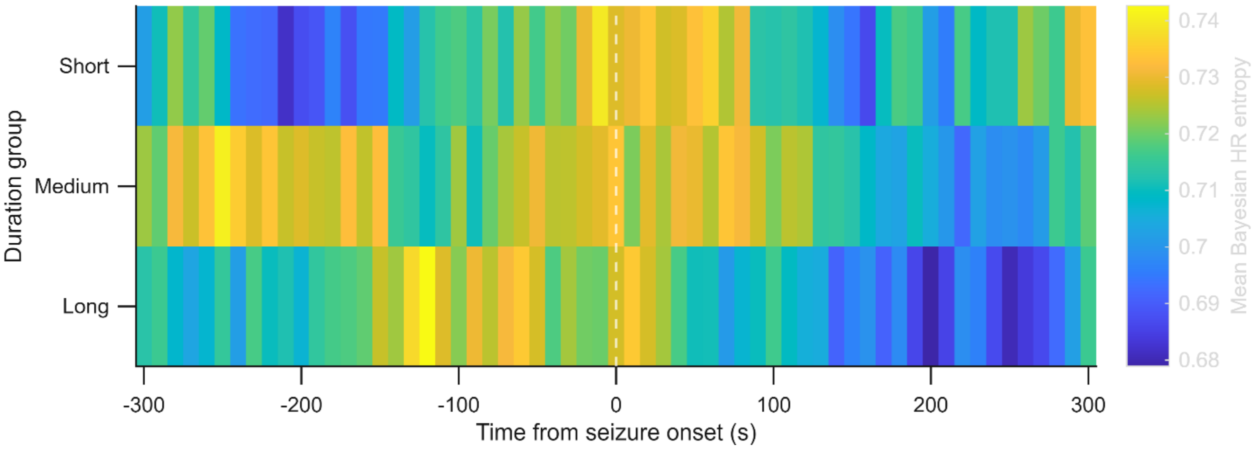


**Supplementary Figure 1 | Mean Bayesian HR entropy trajectories stratified by seizure duration.** Heatmap showing the mean Bayesian HR entropy trajectory for short-, medium-, and long-duration seizures aligned to electrographic seizure onset. Color intensity represents mean Bayesian HR entropy. The vertical dashed line denotes seizure onset (0 s). The heatmap provides a complementary visual representation of the progressive entropy reduction observed in longer seizures.
